# PIQLE: protein-protein interface quality estimation by deep graph learning of multimeric interaction geometries

**DOI:** 10.1101/2023.02.14.528528

**Authors:** Md Hossain Shuvo, Mohimenul Karim, Rahmatullah Roche, Debswapna Bhattacharya

## Abstract

Accurate modeling of protein-protein interaction interface is essential for high-quality protein complex structure prediction. Existing approaches for estimating the quality of a predicted protein complex structural model utilize only the physicochemical properties or energetic contributions of the interacting atoms, ignoring evolutionarily information or inter-atomic multimeric geometries, including interaction distance and orientations. Here we present PIQLE, a deep graph learning method for protein-protein interface quality estimation. PIQLE leverages multimeric interaction geometries and evolutionarily information along with sequence- and structure-derived features to estimate the quality of the individual interactions between the interfacial residues using a multihead graph attention network and then probabilistically combines the estimated quality of the interfacial residues for scoring the overall interface. Experimental results show that PIQLE consistently outperforms existing state-of-the-art methods on multiple independent test datasets across a wide range of evaluation metrics. Our ablation study reveals that the performance gains are connected to the effectiveness of the multihead graph attention network in leveraging multimeric interaction geometries and evolutionary information along with other sequence- and structure-derived features adopted in PIQLE. An open-source software implementation of PIQLE, licensed under the GNU General Public License v3, is freely available at https://github.com/Bhattacharya-Lab/PIQLE.

## Introduction

Protein-protein interactions are the actuators of numerous biological processes (Peng *et al*., 2017). Despite the remarkable progress in predicting single-chain protein structures with very high degree of accuracy (Jumper *et al*., 2021; Baek *et al*., 2021; Wallner, 2022), modeling the structures of protein complexes remains challenging (Zahiri *et al*., 2020; Bryant *et al*., 2022; Evans *et al*., 2022). Traditional protein-protein docking approaches as well as recent deep learning-based protein complex structure prediction methods typically generate a number of candidate structural models and rank them based on estimated confidence scores to select the top-ranked model (Christoffer *et al*., 2021; Pierce *et al*., 2014; Lyskov and Gray, 2008; Gao *et al*., 2022; Bryant *et al*., 2022). With the state-of-the-art protein structure prediction methods approaching near-experimental accuracy on single-chain predictions, accurately modeling the protein-protein interaction interfaces is the key to successfully predicting the structures of protein complexes. As such, high-fidelity estimation of the modeling quality of protein-protein interaction interface from a computationally predicted complex structure is critically important for characterizing protein-protein interactions (Sandor and Kozakov, 2013; Cao and Shen, 2020).

Encouraging progress has been made in protein complex scoring and quality estimation. Physics-based approaches such as ZRANK (Pierce and Weng, 2007) demonstrate effective scoring performance using the weighted sum of several energy terms including van der Waals force, hydrogen bonding, electrostatics, pair potentials, and solvation. ZRANK2 (Pierce and Weng, 2008) further improves the scoring performance by optimizing certain energy terms used in ZRANK. In addition to physics-based approaches, state-of-the-art methods apply machine learning for the quality estimation of complex models. For example, TRScore (Guo *et al*., 2022) estimates the quality protein complex models by learning from a voxelized 3D grid representation of the protein-protein interface using a deep convolutional RepVGG architecture. DOVE (X. Wang *et al*., 2020) applies a 3D convolutional neural network with voxelized representation of protein complexes, while incorporating atomic interaction types and their energetic contributions. Additionally, it integrates knowledge-based statistical potentials GOAP (Zhou and Skolnick, 2011) and ITScore (Huang and Zou, 2008) to capture atomic interaction energies, demonstrating competitive scoring performance. Recently, representation learning with graph neural networks (Zhou *et al*., 2020) is gaining significant attention, leading to the development of several protein complex model quality estimation methods. For example, GNN-DOVE (Wang *et al*., 2021) uses a graph attention network (Veličković *et al*., 2018) by embedding protein complex interfaces as graphs. DPROQ (Chen *et al*., 2022) uses a gated-graph transformer model (GTN) for complex quality estimation.

Despite the progress, these methods do not consider two key factors that can significantly improve protein-protein interface quality estimation performance. First, the geometry of the interaction interface often carries key information for protein complexes (Ganea *et al*., 2022; Dai and Bailey-Kellogg, 2021), but none of the protein complex model scoring methods incorporate multimeric interaction geometries, including the inter-atomic distance and orientations of the residues at the interaction interface. Second, most of the protein complex model scoring approaches typically rely on the physicochemical properties or energetic contributions of the interacting atoms but do not consider the availability of evolutionarily information in the form of multiple sequence alignments (MSAs). That is, they ignore the quality of MSAs during scoring.

Here we present a protein-protein interface quality estimation method called PIQLE by deep graph learning of multimeric interaction geometries. PIQLE formulates protein-protein interface quality estimation as a graph learning task by constructing a graph considering the residues at the interaction interface and estimates the interface quality by training a multihead graph attention network using sequence- and structure-derived node features along with evolutionarily information and newly-introduced edge features in the form of inter-atomic interaction distance and orientations capturing multimeric interaction geometries. Unlike the existing graph neural network-based methods operating on voxelized representation of the protein-protein interface to estimate the overall interface quality, PIQLE first estimates the quality of the individual interactions between the interfacial residues by edge-level error regression and then probabilistically combines the estimated quality of the interfacial residues for scoring the overall interface. Large-scale benchmarking on multiple widely used protein docking decoy sets demonstrates that PIQLE consistently attains better performance than existing complex model quality estimation methods in terms of various evaluation measures including hit rate, success rate, reproducibility of model-native similarity scores, and distinguishability between acceptable and incorrect models. By conducting ablation study on an independent dataset, we directly verify that the improved performance of our method is connected to the effectiveness of the multihead graph attention network in leveraging multimeric interaction geometries and evolutionary information along with the other sequence- and structure-derived features. PIQLE is freely available at https://github.com/Bhattacharya-Lab/PIQLE.

## Materials and methods

**Fig. 1** illustrates our protein-protein interface quality estimation framework consisting of a graph representation of the interaction interface, featurization including multimeric interaction geometries, and quality estimation of the individual interacting residues by edge-level error regression using multihead graph attention network followed by probabilistic combination for the estimation of the overall interface quality.

**Fig. 1.**
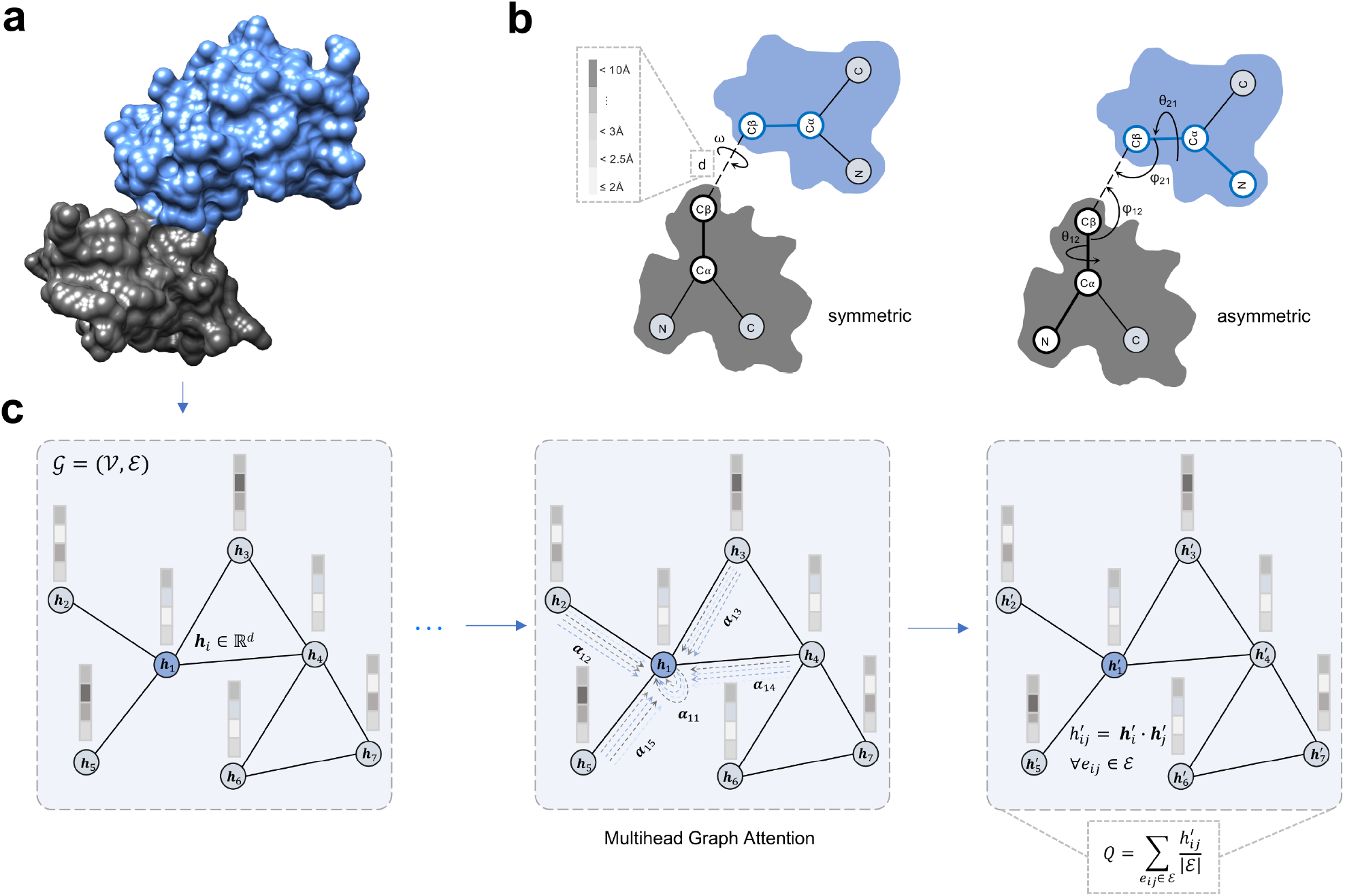
Illustration of the PIQLE framework for protein-protein interface quality estimation.(**a**) The predicted protein complex structure with its two interacting monomers colored in grey and blue. (**b**) Multimeric interaction geometries characterized by the inter-atomic distance and orientations of the residues at the interaction interface. (**c**) Graph representation of the interaction interface and quality estimation of individual interacting residues by edge-level error regression using multihead graph attention network followed by a probabilistic combination.

### Graph representation and featurization

We represent the protein-protein interface as a graph *G* = (*V, E*), in which a node *v* ∈ *V* represents an interface residue and an edge *e∈ E* represents an interacting interface residue pair. We consider an interface residue pair to be interacting if their C_β_ atoms (C_α_ for glycine) are within 10Å (Marze *et al*., 2018). With such a graph representation, we use a total of 17 node features and 27 edge features describing each interface residue and their interactions including sequence- and structure-based node features and multimeric interaction geometric edge features. We describe them below.

### Node features

#### Residue encoding

We cluster 20 naturally occurring amino acids into 4 classes including hydrophobic, hydrophilic, acidic, and basic (**Supplementary Table S1**). For a given residue belonging to one of these classes, we perform one-hot encoding of the residue using 5 class bins with the last bin reserved for the non-standard amino acids belonging to none of the 4 preceding bins, leading to 5 features for each node in the interfacial graph (i.e., a binary vector of 5 entries).

#### Relative residue positioning

To capture the relative positional information for each residue, we extract 1 feature for each node in the interfacial graph corresponding to each of the amino acid residues in a sequence as follows:

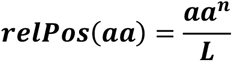

where *aa*^*n*^ is the position of the *n*^*th*^ residue in the sequence and *L* is the length of the sequence.

#### Secondary structure and solvent accessibility

We use DSSP (Kabsch and Sander, 1983) program to calculate the secondary structure and solvent accessibility from the structure. We transform 8-state secondary structures into 3-state by grouping them into helices, strands, and coils for each of the residues in the sequence (**Supplementary Table S1**). Additionally, we discretize the real-valued solvent accessibility into two states of buried and exposed using the solvent-accessible surface area for the corresponding residue (**Supplementary Table S1**). We then use the one-hot encoding of 3-state secondary structures and 2-state solvent accessibility, resulting in 5 features.

#### Local backbone geometry

We calculate phi (*ϕ*) and psi (*ψ*) backbone torsion angles from the structure to capture the local backbone geometry of each residue. We perform sinusoidal and cosine transformations of the angles (Li *et al*., 2017), leading to 4 features.

#### Evolutionarily information

We compute the number of effective sequences (*N*_*eff*_) from the MSAs of the individual monomer and concatenated MSA of the complex to account for the depth of the MSAs, thereby considering the availability of evolutionarily information. To generate MSAs from an individual monomeric sequence, we run HHBlits (Remmert *et al*., 2012) for three iterations with an E-value inclusion threshold of 10^3^ for searching against database Uniclust30 (Mirdita *et al*., 2017) database with a query sequence coverage of 10% and pairwise sequence identity of 90%. We then calculate the normalized number of sequences as:

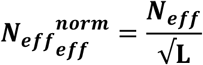

where *N*_*eff*_ is the reciprocated sum of the number of sequences in the MSA having a sequence identity greater than 80% to the *n*^*th*^ sequence and *L* is the length of the sequence (Li *et al*., 2019). Additionally, we concatenate both of the MSAs of the individual monomeric sequences using GLINTER (Xie and Xu, 2022) and compute *N*_*eff*_ of the concatenated MSA using the same aforementioned strategy. The *N*_*eff*_ computed from the MSAs of the individual monomers and the concatenated MSA result in 2 features.

### Edge features

#### Multimeric interaction distance

To capture multimeric interaction geometry, we discretize the Euclidian distance between the C_β_ atoms (C_α_ for glycine) of the interacting interface residue pairs into 17 bins ranging from 2Å to 10Å having a bin width of 0.5Å. The discretized interaction distance is represented by one-hot encoding, resulting in 17 edge features.

#### Multimeric interaction orientation

In addition to C_β_-C_β_ distances, we also include the orientations of the interacting interface residue pairs by extending the work of trRosetta (Yang *et al*., 2020) for multimers. In particular, our multimeric interaction orientation is represented by 3 torsion (ω, θ_12_, θ_21_) and 2 planar angles (ϕ_12_, ϕ_21_), as shown in **Fig. 1b**. The ω torsion angle measures the rotation along the virtual axis connecting the C_β_ atoms of the interacting interface residue pairs, and θ_12_, ϕ_12_ (θ_21_, ϕ_21_) angles specify the direction of the C_β_ atom of interface residue of the first (second) interacting monomer in a reference frame centered on the interface residue of the second (first) interacting monomer. Unlike the symmetric torsion angle ω, θ and ϕ are asymmetric and depend on the order of the monomeric interacting interface residue pairs. Once again, we perform sinusoidal and cosine transformations of the angles, leading to 10 features.

##### Network architecture

**Fig. 1c** shows the architecture of our multihead graph attention network for protein-protein interface quality estimation. The network consists of 4 multihead graph attention layers (Veličković *et al*., 2018). All the intermediate layers have four attention heads except for the output layer, which has one attention head. We perfom hyperparameter selection on an independent validation set using grid search to determine the optimal number of layers and heads (**Supplementary Table S2**). The input layer of the network takes the interfacial graph *G* consisting of nodes *V* and edges *E* with the associated nodes and edge features as *G*(*V*_*i*_ ∈ ℝ^ν×0^ × *E*_*ij*_ ∈ ℝ^*e*×1^). We use an empirically selected hidden dimension of 32 for the input layer with a scaling factor of 0.5 for each succeeding layers. Additionally, we perform a concatenation operation of all the heads along the output dimension of 1. Therefore, the output dimension of each intermediate layer depends on the number of hidden dimensions, and thus the output dimension of the multihead attention layer *l* is as follows:

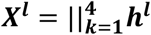

where, *k* represents the number of heads and *h* is the hidden dimension at layer *l*. Of note, each multihead attention layer performs a series of operations before it feeds the output to the next layer. First, we concatenate both the node and edge features and perform a linear transformation to embed both the node 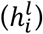 and edge 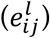 input features with the initialized weight *W* to d-dimensional hidden features assigned to each node (Veličković *et al*., 2018; Dwivedi *et al*., 2022) as follows:

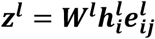

where, *z* represents the embedded features at layer *l* and *W* is the learnable network parameters, normalized using the Xavier weight initialization procedure at each layer to prevent vanishing and exploding gradient problems (Glorot and Bengio, 2010). We then compute an attention score *a*_*ij*_ between the neighboring nodes of each edge by performing self-attention on the incident nodes as:

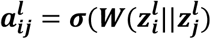

where, 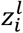 and 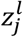 are the embeddings of the incident nodes of an edge. Both embeddings are concatenated, and a dot product is computed with a learnable weight vector *W* (*W* ∈ ℝ^**D**^) where *D* represents the input dimension. Meanwhile, the node features of each node are updated with the combination of neighboring node features and the attention score *a*_*ij*_ as follows:

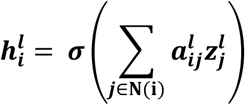

##### Model training

For the assignment of the ground truth interface quality score during training, we first calculate the observed C_β_-C_β_ distance between the interacting interface residue pairs in the predicted complex structural model 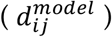 and the corresponding residue pairs in the native structure 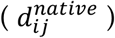. We then assign a normalized ground truth interface quality score *z*_*ij*_ to the edge *e*_*ij*_ as follows:

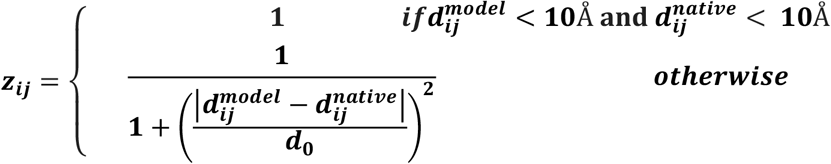

where, 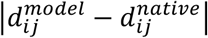 is the observed edge-level error between the interacting interface residue pairs corresponding to the edge *e*_*ij*_, and *d*_0_ is a normalizing constant whose value is set to 10Å.

During model training, we learn the interface quality score *z*_*ij*_ for each edge *e*_*ij*_ through edge-level error regression by optimizing the Mean Squared Error (MSE) loss function with sum reduction using the Deep Graph Library (DGL) (M. Wang *et al*., 2020). We use the Adam optimizer (Kingma and Ba, 2014) with a learning rate of 0.001 and a weight decay of 0.0005. The training process consists of at most 500 epochs on an NVIDIA A40 GPU having an early stopping criterion with patience set to 40 to prevent overfitting.

##### Estimation of protein-protein interface quality

During the inference, we first estimate the interface quality score for each of the interacting residue pairs in the interface graph through edge-level error regression by computing the dot product between the predicted embeddings of the corresponding nodes as follows:

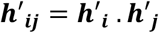

where, *h*′_*i*_ and *h*′_*j*_ represent the node embeddings of nodes *i* and *j* connected by the edge *e*_*ij*_ in the final layer of the multi-head graph attention network. We then probabilistically combine the estimated quality scores of the individual interfacial residue pairs for estimating the overall interface quality score *Q* as follows:

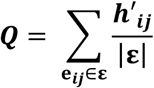

where, |ε| represents the number of interfacial residue pairs in the model and *h*′_*ij*_ is the estimated interaction score for the edge *e*_*ij*_. The overall interface quality score *Q* ranges between 0 and 1 with a higher score indicating better protein-protein interface quality.

##### Datasets

To train the graph attention network of PIQLE, we use the docking benchmark set of Dockground (Kundrotas *et al*., 2018) version 2 (hereafter called Dockground v2) containing 179 dimeric protein complex targets having the length ranging from 92 to 894 residues with 100 complex structural models for each target, generated by docking the unbound structure of the receptor to the ligand.

To benchmark our method, we use the docking benchmark set of Dockground version 1 (hereafter called Dockground v1), comprising of 61 dimeric protein complex targets having the length ranging from 107 and 892 residues. We discard all targets from the Dockground v1 test dataset overlapping with our training set Dockground v2 using an average pairwise sequence identity cutoff of 20%, resulting in 23 dimer targets with an average of 109 decoys per target. Additionally, we use the Heterodimer-AF2 (hereafter called HAF2) (Chen *et al*., 2022) dataset consisting of 13 targets having an average of 105 decoys per target with the length ranging from 78 to 1248 residues generated using AlphaFold-Multimer (Evans *et al*., 2022).

For ablation studies, we use the docking Benchmark version 4.0 (Hwang *et al*., 2010) comprising of 69 dimeric protein complex targets having the length ranging from 23 to 822 residues that are non-overlapping to the targets in the training and benchmarking datasets (pairwise sequence identity cutoff of 20%) with each target having 100 complex structural models generated by ZDOCK (Pierce *et al*., 2014). It is worth noting that all the datasets used for training, benchmarking, and ablation studies are non-overlapping with an average pairwise sequence identity of less than 20% between any pair of datasets (**Supplementary Table S3**).

##### Evaluation metrics and competing methods

We assess the performance of our method using various evaluation metrics based on the DockQ score (Basu and Wallner, 2016) as the ground truth. We use three evaluation criteria to measure the protein-protein interface quality estimation performance: (i) ability to reproduce the ground truth DockQ scores, (ii) ability to rank complex structural models, and (iii) ability to distinguish acceptable from incorrect models. For the first criterion, we use the Spearman correlation coefficient (ρ) between the estimated quality of the protein complexes and their corresponding DockQ scores. Consequently, a higher correlation indicates better reproducibility. For the second criterion, we use the top-N success rate (herein: *SR*) and top-N hit rate (herein: *HR*) (Guo *et al*., 2022). The top-N success rate is calculated as the percentage of complex targets having at least one acceptable model among top N ranked models as follows:

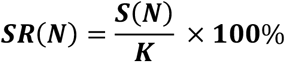

where, *S*(*N*) is the number of complex targets having at least one acceptable model among top N ranked models and *K* is the total number of targets, where a cutoff of DockQ = 0.23 is used to identify acceptable models (Basu and Wallner, 2016). The top-N hit rate is calculated as the fraction of acceptable models among top-ranked models relative to all acceptable models in the entire dataset as follows:

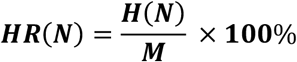

where, *H*(*N*) is the total number of acceptable models among top N ranked models and *M* is the total number of acceptable models in the dataset. Higher success and hit rate indicate better ranking ability, especially for low values of N. We evaluate the success and hit rate of top-ranked models for various values of N including top-1, top-5, top-10, top-15, top-20, top-25, and top-30. For the third criterion, we perform receiver operating characteristics (ROC) analysis using a DockQ score cutoff of 0.23 to separate acceptable and incorrect models. Meanwhile, the Area Under the ROC curve (AUC) quantifies the ability of a method to distinguish between acceptable and incorrect models with a higher AUC indicating better distinguishing ability.

We compare the performance of PIQLE against a number of existing protein complex quality estimation methods ranging from physics-based approaches to machine learning-based methods. As a representative physics-based method, we compare PIQLE against ZRANK2 (Pierce and Weng, 2008), which is an improved version of ZRANK (Pierce and Weng, 2007). For fair comparison, we use min-max normalization strategy to scale ZRANK2 energy scores to the same range as the predicted scores of other methods including PIQLE as follows:

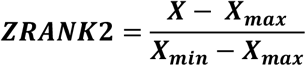

where, X is the raw ZRANK2 estimated energy scores and *X*_*min*_ and *X*_*max*_ represent the smallest and largest estimated scores, respectively, considering all predicted complex structural models for a specific target.

We also compare the performance of PIQLE against various machine learning-based approaches including 3D Convolutional Neural Network (3DCNN)-based methods TRScore (Guo *et al*., 2022) and four variants of DOVE (X. Wang *et al*., 2020): DOVE-Atom20, DOVE-Atom40, DOVE-GOAP, and DOVE-Atom40+GOAP as well as recent graph neural network-based methods GNN-DOVE (Wang *et al*., 2021) and DPROQ (Chen *et al*., 2022). We exclude the comparison to GNN-DOVE and TRScore on the Dockgorund v1 test dataset due to the overlap of the training datasets used in GNN-DOVE and TRScore with the complex targets present in the Dockground v1 dataset. Meanwhile, all methods are included for performance comparison on the HAF2 test dataset.

## Results

### Reproducing ground truth DockQ scores

**Fig. 2** shows the Spearman correlation coefficients (ρ) between the estimated qualities of the protein-protein interfaces and their corresponding ground truth DockQ scores for PIQLE and the other competing methods. PIQLE consistently outperforms all other competing methods in both Dockground v1 and HAF2 datasets. On Dockground v1, PIQLE attains the highest Spearman correlation of 0.519 which is much better than the second-best 3D Convolutional Neural Network (3DCNN)-based method DOVE-Atom20 (0.305), the recent graph transformer network (GTN)-based method DPROQ (0.103), and physics-based scoring function ZRANK2 (0.159). The same trend continues for the HAF2 dataset, in which PIQLE attains the highest Spearman correlation of 0.429 that is significantly better than the other competing methods. 3DCNN -based method DOVE remains as the second-best method with its variant DOVE-Atom+GOAP attaining a Spearman correlation of 0.382. The recent graph neural network-based methods DPROQ and GNN-DOVE, however, fail to generalize on the HAF2 dataset attaining negative Spearman correlations of -0.014 and -0.331, respectively. In summary, our method PIQLE delivers improved ability to reproduce ground truth DockQ scores with high fidelity.

**Fig. 2.**
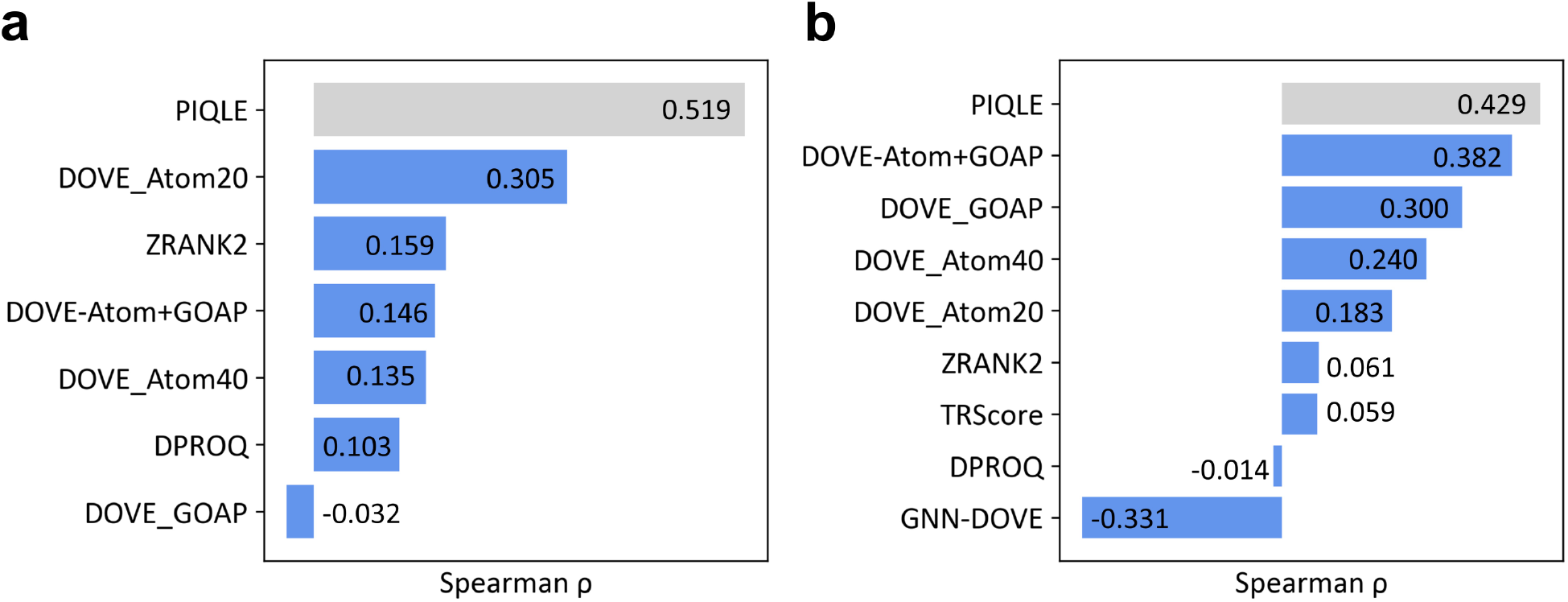
Reproducibility of ground truth DockQ scores for PIQLE (in gray) and the competing methods (blue), sorted in decreasing order of Spearman correlations coefficient (ρ) between the estimated qualities of the protein-protein interfaces and their corresponding DockQ scores on (**a**) Dockground v1 and (**b**) HAF2 datasets.

### Ranking complex structural models

**Fig. 3a-3b** show the complex model ranking performance of PIQLE and the other competing methods in terms of the success rate (*SR*) metric, which evaluates the ability of a method to select at least one acceptable model within top N ranked models. As shown in **Fig. 3a**, PIQLE consistently achieves the highest *SR* among all methods for almost all top-N rankings in the Dockground v1 dataset. The noticeably higher *SR* of PIQLE at low values of N such as top-1 (∼44%) and top-5 (∼74%) is particularly noteworthy. In the HAF2 dataset (**Fig. 3b**), PIQLE attains a higher top-1 *SR* of ∼77% compared to the other methods, whereas some of the other methods such as ZRANK2 and DOVE-ATOM20 achieve comparable or higher *SR* values, particularly for high values of N. Overall, PIQLE frequently attains higher success rates, particularly when N is low.

**Fig. 3.**
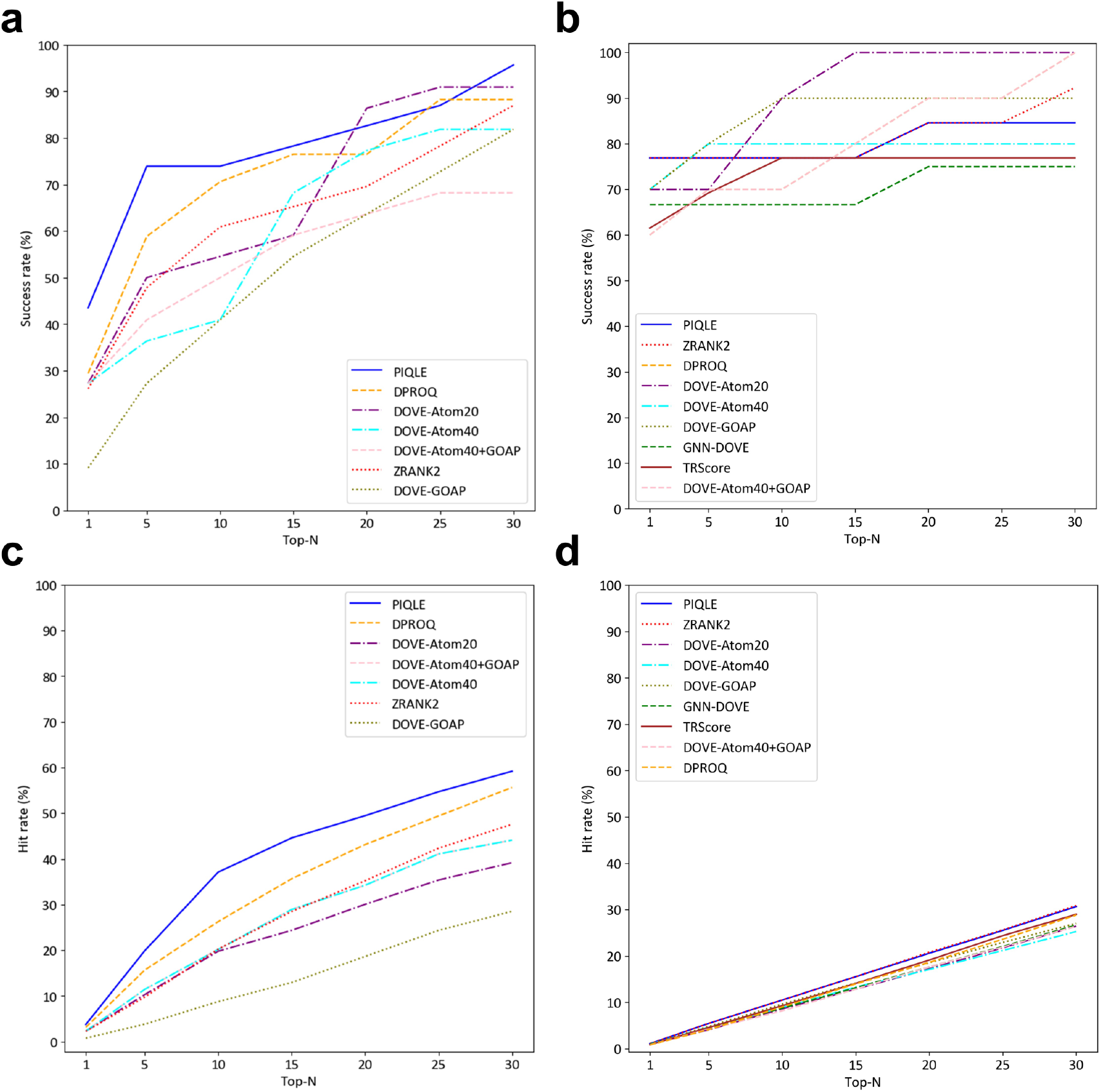
Ranking complex structural models for PIQLE and the competing methods in terms of success rate on (**a**) Dockground v1 dataset, (**b**) HAF2 dataset and hit rate on (**c**) Dockground v1 dataset, (**d**) HAF2 dataset based on top-1, top-5, top-10, top-15, top-20, top-25, and top-30 models. A cutoff of DockQ = 0.23 is used to identify acceptable models.

**Fig. 3c-3d** show the ranking ability of PIQLE and the other competing methods in terms of the hit rate (*HR*) metric, which evaluates the performance of a method based on the total number of acceptable models among top-ranked models relative to all acceptable models in the entire dataset. As shown in **Fig. 3c**, PIQLE significantly outperforms all other competing methods by achieving the highest *HR* on Dockground v1 dataset, for all values of N. For example, PIQLE improves the top-10 *HR* by more than 40% (37.079 vs 26.250) over the second-best method DPROQ. PIQLE also consistently attains better *HR* performance on the HAF2 dataset as shown in **Fig. 3d**. It is interesting to note the somewhat low *HR* performance of all methods including ours on the HAF2 dataset. While all methods, particularly the machine learning-based approaches achieve higher *SR* on the HAF2 dataset, they appear to be less effective at selecting a large proportion of acceptable models from a smaller number of top-ranked models measured by *HR*, suggesting a need for further improvement. Nevertheless, our new method PIQLE strikes an ideal balance to deliver top performance in terms of both success rate and hit rate across different datasets, indicating its all-round ability in ranking complex models.

### Distinguishing acceptable from incorrect models

In addition to reproducing the ground truth DockQ scores with high fidelity and accurately ranking complex structural models, the ability to distinguish acceptable from non-acceptable prediction is critically important. **Fig. 4** shows the area under ROC curve (AUC) attained by PIQLE and the competing methods on the test datasets. PIQLE consistently outperforms all other competing methods by achieving the best AUC on both Dockground v1 and HAF2 dataset. For the Dockground v1 dataset, AUC attained by PIQLE is 0.711 which is closely followed by the second-best performing method DOVE-Atom40+GOAP with an AUC of 0.710. On the HAF2 dataset, PIQLE attains an AUC of 0. 743, which is significantly higher than all competing methods including the second-best method DOVE-Atom40+GOAP having an AUC of 0.648. In summary, PIQLE exhibits an improved ability to distinguish between acceptable and non-acceptable prediction.

**Fig. 4.**
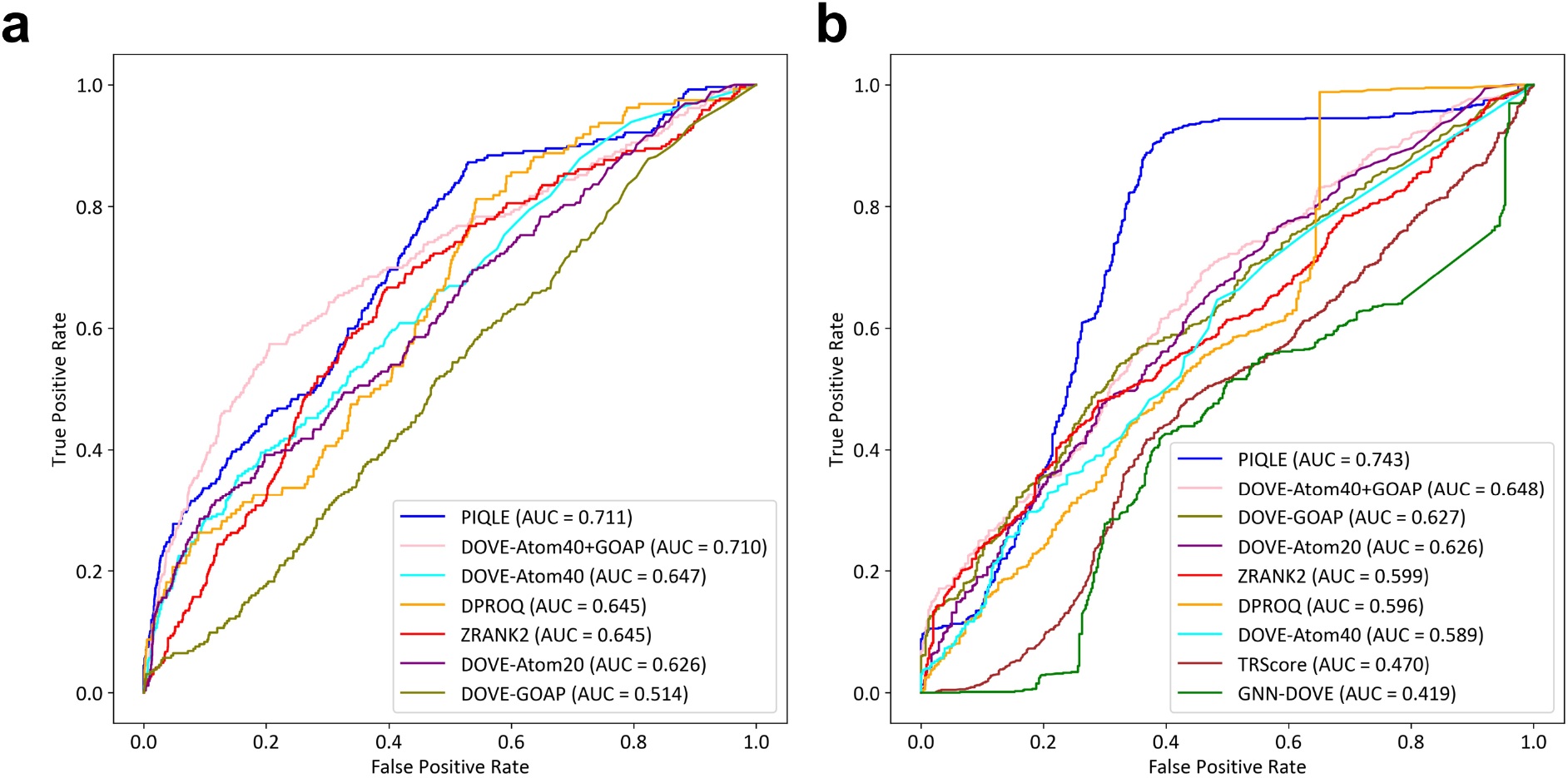
Distinguishability of acceptable vs. incorrect models for PIQLE and the competing methods on (**a**) Dockground v1 and (**b**) HAF2 datasets. A cutoff of DockQ = 0.23 is used to separate acceptable from incorrect models.

### Case study

**Fig. 5** shows some representative examples of protein-protein interface quality estimation by PIQLE for selected targets from Dockground v1 and HAF2 datasets with varying degree of predictive modeling accuracy. For a reasonably well predicted complex structural model for Dockground v1 target 1r0r having a DockQ score of 0.732 (**Fig. 5a**), PIQLE estimates an interfacial quality score of 0.625. Apart from a few false positive interacting residue pairs, most of the interface regions in this predicted complex structural model are correct with an F1 score of 0.674 considering the previously defined C_β_-C_β_ distance threshold of 10Å (Marze et al., 2018) for identifying the true interacting residue pairs. For a moderate-quality predicted complex structural model for HAF2 target 7nkz having a DockQ score of 0.478 and several false positive interacting residue pairs with an F1 score of 0.333 (**Fig. 5b**), PIQLE estimates a moderate interfacial quality score of 0.398. Additionally, **Fig. 5c-d** show two low-quality predicted complex structural models for Dockground v1 target 1ppf and HAF2 target 7lxt having DockQ score of 0.102 and 0.127, respectively, with noticeably wrong interfaces. For these models, PIQLE estimates much lower interfacial quality scores of 0.143 and 0.130, respectively.

**Fig. 5.**
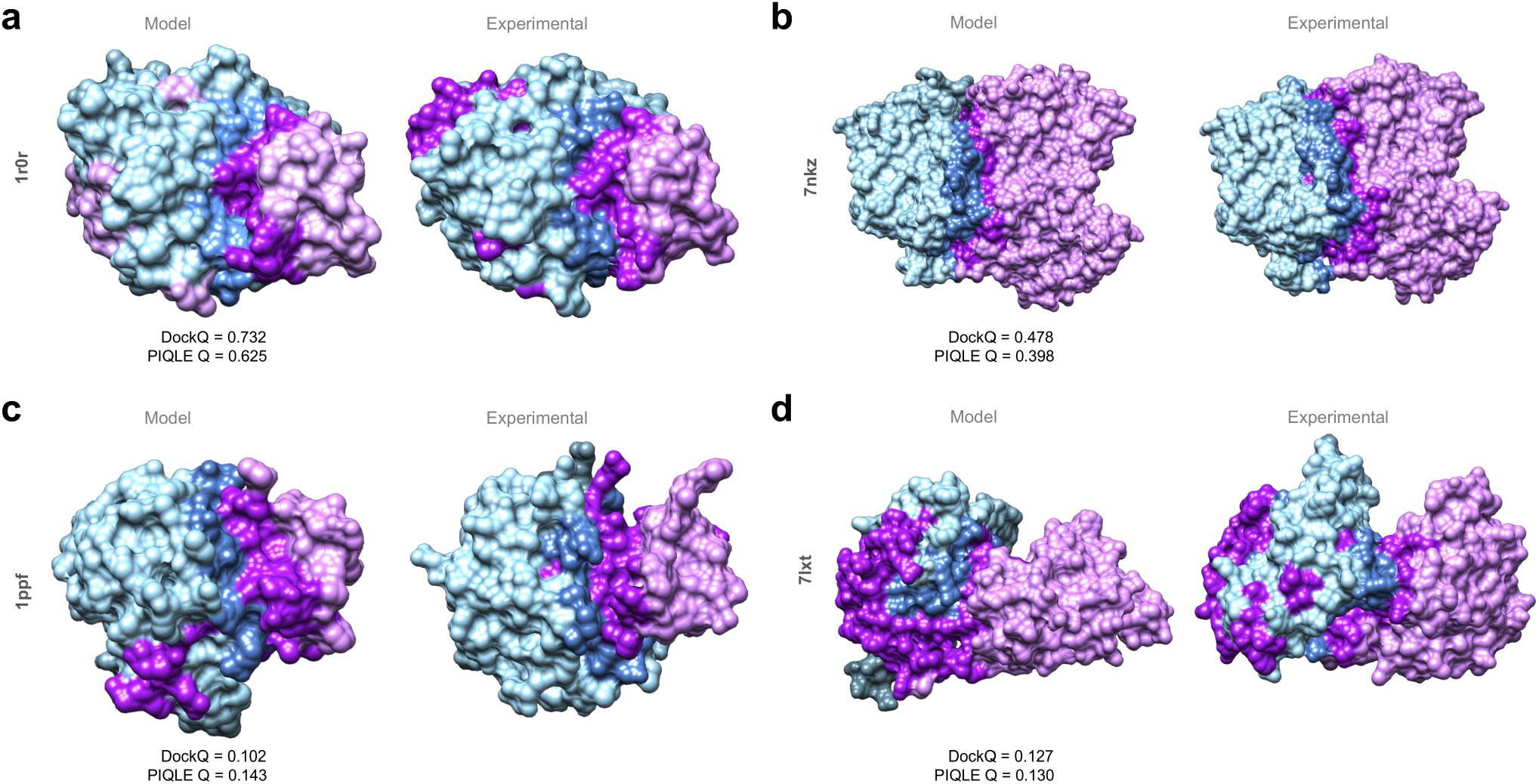
Case study on protein-protein interface quality estimation by PIQLE using predicted complex structural models for (**a**) Dockground v1 target 1r0r, (**b**) HAF2 target 7nkz, (**c**) Dockground v1 target 1ppf, (**d**) HAF2 target 7lxt. For each target, the interacting protein chains are colored in blue (chain 1) and purple (chain 2) with the interface regions highlighted in darker shades of blue and purple. The experimental structures for each target are shown side-by-side with the observed interface regions annotated.

### Ablation study

To examine the relative importance of the features adopted in PIQLE, we conduct feature ablation experiments by gradually isolating the contribution of individual feature or groups of features during model training and evaluating the accuracy on the independent ZDOCK validation dataset. **Fig. 6a** shows the Spearman correlation coefficients (ρ) between the estimated qualities of the protein-protein interfaces and their corresponding ground truth DockQ scores when various features are isolated from the full-fledged version of PIQLE. The results demonstrate that all features contribute to the overall performance achieved by PIQLE. For example, we notice an accuracy decline when we isolate the sequence-based features one by one including amino acid residue encoding (No residue encoding) and relative residue positioning (No relative residue positioning). Importantly, discarding the evolutionarily information noticeably declines the overall performance (No evolutionarily information), indicating the effectiveness of MSA-derived evolutionarily information. Not surprisingly, we notice a dramatic performance drop when both the residue-based features and the evolutionary information are isolated (No residue + evolutionarily information). Similarly, we also notice a performance drop when the feature based on local backbone geometry is discarded (No local backbone geometry). Additionally, we notice a consistent accuracy decline when we discard the newly-introduced edge features based on multimeric interaction distance (No multimeric interaction distance), multimeric orientation (No multimeric interaction orientation) and their combination (No multimeric interaction geometry). That is, the improved performance of our method is connected to the effective integration of multimeric interaction geometries.

**Fig. 6.**
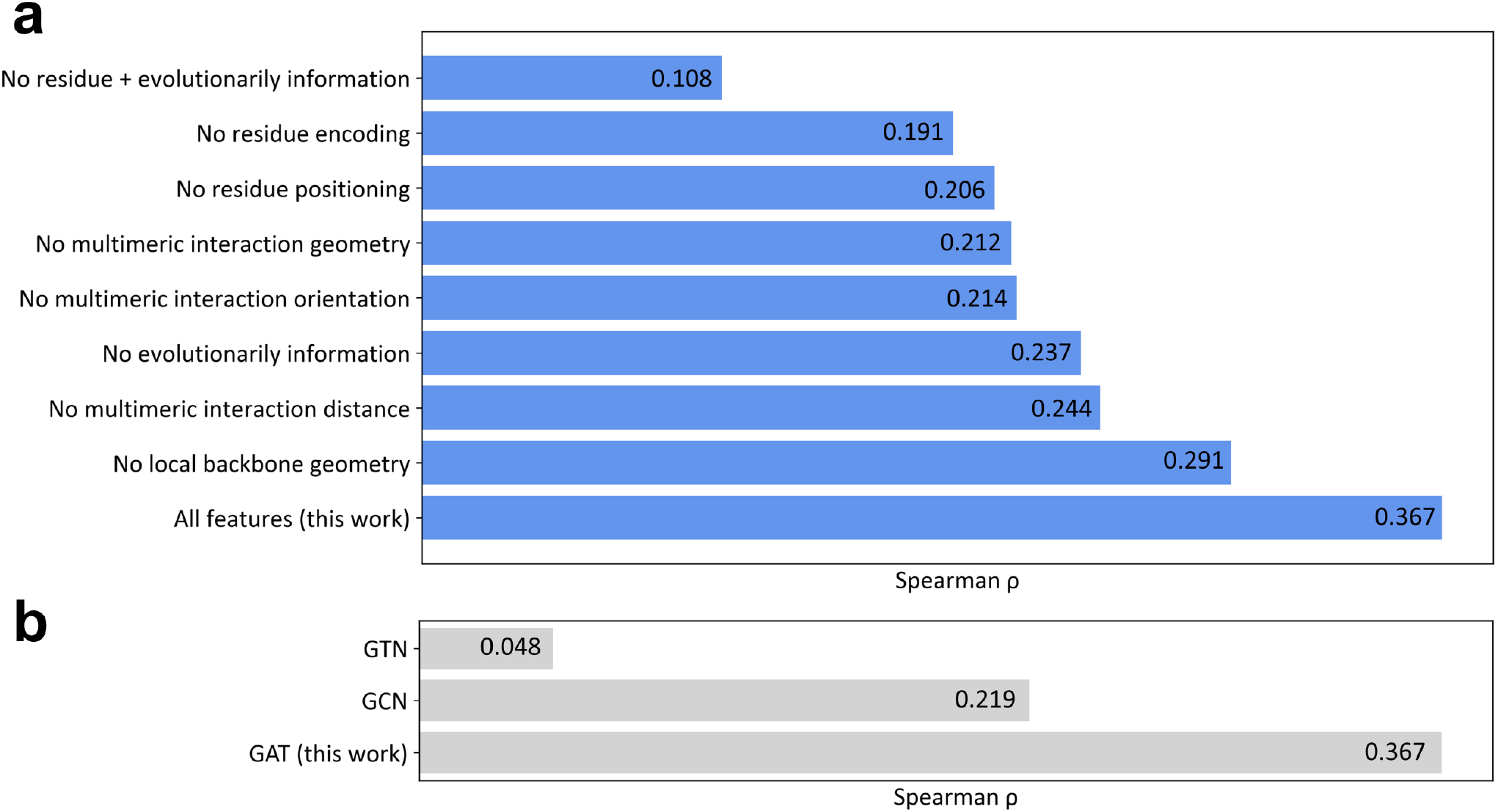
Ablation study on the independent ZDOCK validation dataset in terms of Spearman correlations coefficient (ρ) between the estimated qualities of the protein-protein interfaces and their corresponding DockQ scores by (**a**) gradually isolating individual feature or groups of features during model training, and (**b**) training two baseline graph neural network models employing graph convolutional network (GCN) and graph transformer network (GTN) architectures.

To further investigate the contribution of the multi-head graph attention network model used in PIQLE, we train two baseline graph neural network (GNN)-based models for protein-protein interface quality estimation: graph convolutional network (GCN) (Kipf and Welling, 2017) and graph transformer network (GTN) (Yun *et al*., 2019). All baseline networks are trained on the same training dataset using the same set of input features and hyperparameters as the full-fledged version of PIQLE. **Fig. 6b** shows the performance of PIQLE compared to the baseline networks on the independent ZDOCK validation dataset in terms of the Spearman correlation coefficients (ρ) between the estimated qualities of the protein-protein interfaces and their corresponding ground truth DockQ scores. The multihead graph attention network architecture of PIQLE significantly outperforms the other baseline networks, demonstrating its effectiveness for protein-protein interface quality estimation task.

## Conclusion

This work introduces PIQLE, a new method for protein-protein interface quality estimation by deep graph learning of multimeric interaction geometries. PIQLE exploits multihead graph attention network architecture leveraging multimeric interaction geometries and evolutionarily information along with sequence- and structure-derived features to estimate the quality of the individual interactions between the interfacial residues and then probabilistically combines the estimated quality of the interfacial residues for scoring the overall interface. We demonstrate that PIQLE attains state-of-the-art protein-protein interface quality estimation performance by conducting large-scale benchmarking on multiple widely used protein docking decoy sets. Our ablation study on an independent validation set confirms the contribution of various features adopted in PIQLE and the effectiveness of the multihead graph attention network architecture.

Our study leads to a number of future directions to consider: of particular interest is the possibility of broadening the applicability of our method for higher order oligomers and large protein assemblies. Further, a promising direction for future work is to consider the diversity of predictive modeling ensemble and conformational states of the interacting monomers for interface quality estimation for interacting proteins having multi-state conformational dynamics. Finally, integrating complementary features such as residue-level self-assessment confidence estimates for the interacting protein chains and sequence-based disorder prediction coupled with richer deep graph representation learning framework may further boost protein-protein interface quality estimation performance. We expect our method can be extended to other biomolecular interface characterization, including estimating the quality of predicted protein interaction with other molecules, such as DNA, RNA, and small ligands.

## Data availability

The raw data used in this study, including the datasets for train, test, and validation are collected from publicly available sources. The Dockground v2 training set is available at https://dockground.compbio.ku.edu/downloads/unbound/decoy/decoys-set2-1.0.tgz. The Dockground v1 test set is available at https://dockground.compbio.ku.edu/downloads/unbound/decoy/decoys1.0.zip. The Heterodimer-AlphaFold2 test set is available at https://zenodo.org/record/6569837/files/DproQ_benchmark.tgz. The ZDOCK docking benchmark version 4.0 validation set is available at http://zlab.umassmed.edu/benchmark/.

## Code availability

An open-source software implementation of PIQLE, licensed under the GNU General Public License v3, is freely available at https://github.com/Bhattacharya-Lab/PIQLE.

## Acknowledgements

This work was partially supported by the National Institute of General Medical Sciences [R35GM138146 to D.B.] and the National Science Foundation [DBI2208679 to D.B.].

