## Supplementary Information for "PIQLE: protein-protein interface quality estimation by deep graph learning of multimeric interaction geometries"

*for*

**Supplementary Table S1.** Categorization of several node features adopted in PIQLE.

| | **Properties** | **Category** |
|---|---|---|
| 5-state amino acid residue types | ALA VAL LEU ILE PRO PHE MET TRP | Hydrophobic |
| | GLY SER THR CYS TYR ASN GLN | Hydrophilic |
| | LYS ARG HIS | Basic |
| | ASP GLU | Acidic |
| | Any other residues | Neutral/None |
| 3-state secondary structure types | G, H, I | H |
| | E, B | E |
| | Other | C |
| 2-state solvent accessibility types | RSA (ACC / Max. SA) < 25% | B |
| | RSA (ACC / Max. SA) > 25% | E |

**Supplementary Table S2.** Hyperparameter search for graph attention network on the independent ZDOCK validation dataset in terms of Spearman correlations coefficient (ρ) between the estimated protein-protein interface quality scores and their corresponding DockQ scores.

| **Hyperparameter** | | **Spearman ρ** |
|---|---|---|
| **# Layers** | **# Heads** | |
| **4** | 2 | 0.168 |
| | **4** | **0.367** |
| | 6 | 0.199 |
| | 8 | 0.266 |
| 6 | 2 | 0.155 |
| | 4 | 0.178 |
| | 6 | 0.300 |
| | 8 | 0.288 |
| 8 | 2 | 0.222 |
| | 4 | 0.280 |
| | 6 | 0.325 |
| | 8 | 0.289 |

| | | |
|---|---|---|
| 10 | 2 | 0.200 |
| | 4 | 0.189 |
| | 6 | 0.172 |
| | 8 | 0.185 |
| 12 | 2 | 0.173 |
| | 4 | 0.218 |
| | 6 | 0.230 |
| | 8 | 0.227 |
| 14 | 2 | 0.220 |
| | 4 | 0.199 |
| | 6 | 0.202 |
| | 8 | 0.235 |
| 16 | 2 | 0.139 |
| | 4 | 0.238 |
| | 6 | -0.020 |
| | 8 | 0.225 |

*Note:* Graph attention network with the hyperparameters in bold yield the highest Spearman ρ

**Supplementary Table S3.** Pairwise sequence identity between training, testing, and validation datasets.

| Datasets | Dockground v1[a] | Dockground v2[b] | ZDOCK[c] | HAF2[d] |
|---|---|---|---|---|
| **Dockground v1** | | 0.183 | 0.192 | 0.162 |
| **Dockground v2** | 0.183 | | 0.179 | 0.152 |
| **ZDOCK** | 0.192 | 0.179 | | 0.133 |
| **HAF2** | 0.162 | 0.152 | 0.133 | |

[a, d]Testing datasets
[b]Training dataset
[c]Validation dataset
[d]Heterodimer-AF2 dataset
